# Symplasmic phloem loading and subcellular transport in storage roots are key factors for carbon allocation in cassava

**DOI:** 10.1101/2024.02.21.581442

**Authors:** David Rüscher, Viktoriya V. Vasina, Jan Knoblauch, Leo Bellin, Benjamin Pommerrenig, Saleh Alseekh, Alisdair R. Fernie, H. Ekkehard Neuhaus, Michael Knoblauch, Uwe Sonnewald, Wolfgang Zierer

## Abstract

Cassava is a deciduous woody perennial shrub that stores large amounts of carbon and water in its storage roots. Previous studies have shown that assimilate unloading into storage roots happens symplasmically once secondary anatomy is established. However, mechanisms controlling phloem loading and overall carbon partitioning to different cassava tissues remain unclear. Here we used a combination of histological, transcriptional, and biochemical analyses on different cassava tissues and timepoints to better understand source-sink carbon allocation. We find that cassava likely utilizes a predominantly passive symplasmic phloem loading strategy, indicated by the lack of expression of genes coding for key players of sucrose transport, the existence of branched plasmodesmata in the companion cell/bundle sheath interface of minor leaf veins, and very high leaf sucrose concentrations. Furthermore, we show that tissue-specific changes in anatomy and NSC contents are associated with tissue-specific modification in gene expression for sucrose cleavage/synthesis, as well as subcellular compartmentalization of sugars. Overall, our data suggest that carbon allocation during storage root filling is mostly facilitated symplasmically, and is likely mostly regulated by local tissue demand and subcellular compartmentalization.

## Introduction

Cassava (*Manihot esculenta*) is an important crop in Sub-Saharan Africa, South America and Southeast Asia. Its starchy storage roots are crucial for food security, especially in Sub-Saharan Africa, where over 60% of the global storage root production is realized (FAOSTAT, 2022). Despite an overall increase in the worldwide cassava production, yields in Sub-Saharan Africa are largely stagnant (FAOSTAT, 2022). This, combined with an increasing population and decrease in the productivity of other crops due to the advancing climate change, makes a sustainable increase in cassava storage root yield paramount for future food security. To this end, advances in cassava breeding and biotechnology are important, which require a better understanding of its physiology.

Cassava is a woody perennial originating from the tropical savanna regions of South America, where it was cultivated over 6000 years ago (Olsen and Schaal, 1999). Perennial shrubs and trees of these areas follow a deciduous life cycle, losing their leaves during the dry season and regrowing them during the rainy season (Opler *et al*., 1976). This growth pattern makes effective storage of carbon and water important. Woody plants can store impressive amounts of non-structural carbohydrates (NSC), such as starch, in their trunks and roots (Loescher et al., 1990, Furze et al., 2018). Cassava, in addition to its stem storage tissues, possesses dedicated storage roots that support the accumulation of large amounts of carbon and water. Through the activity of a single vascular cambium in the storage root, large amounts of storage parenchyma cells are produced and quickly filled with high levels of starch, in addition to high levels of free amino acids, sucrose and hexoses (Rüscher et al., 2021). Previous studies have shown that, similar to other woody species, phloem unloading in the secondary vasculature of cassava occurs largely symplasmically (Mehdi et al., 2019, Pan et al., 2021). However, it is not clear how assimilates are partitioned between leaves, stems, and storage roots, and how this is partitioning is regulated throughout storage root development. However, this process is critical because ineffective carbon flux to storage roots negatively affects yield (Ruiz-Vera et al., 2020, Chiewchankaset et al., 2022). Similarly, the mechanism by which assimilates enter the phloem system in the first place remain unclear to date.

To gain greater insight into the processes regulating carbon allocation and storage in cassava, different cassava tissues and time points were analyzed in terms of their anatomical, transcriptional, and metabolic characteristics. The obtained data indicate that cassava predominantly utilizes a passive symplasmic phloem loading strategy, a conclusion that was further supported using electron microscopy. Furthermore, the importance of local sink strength and intercellular compartmentation of sugars in the storage root is highlighted and supported both by transcriptional analyses and metabolite measurements.

## Results

### Changes in plant anatomy and NSC content during root bulking

Storage roots initially develop from fibrous roots by initiating secondary growth and subsequently produce large amounts of storage parenchyma cells (Chaweewan and Taylor, 2015, Rüscher et al., 2021). This process of lateral size increase and starch filling is referred to as bulking, and is associated with a pronounced accumulation of carbon compounds (Rüscher et al., 2021). To gain further insights into the processes regulating carbon allocation and storage in cassava, we analyzed source leaves, stem tissues, fibrous roots and storage roots at distinct stages of storage root development. Following earlier studies, we chose the timepoint of initial storage root vascular cambium formation (“pre-bulking”; PB), the initial expansion of the root xylem via woody tissue deposition (“early-bulking”; EB), and the stage of storage parenchyma formation (“during bulking”; DB) in the storage root (Fig. 1).

**Fig. 1.**
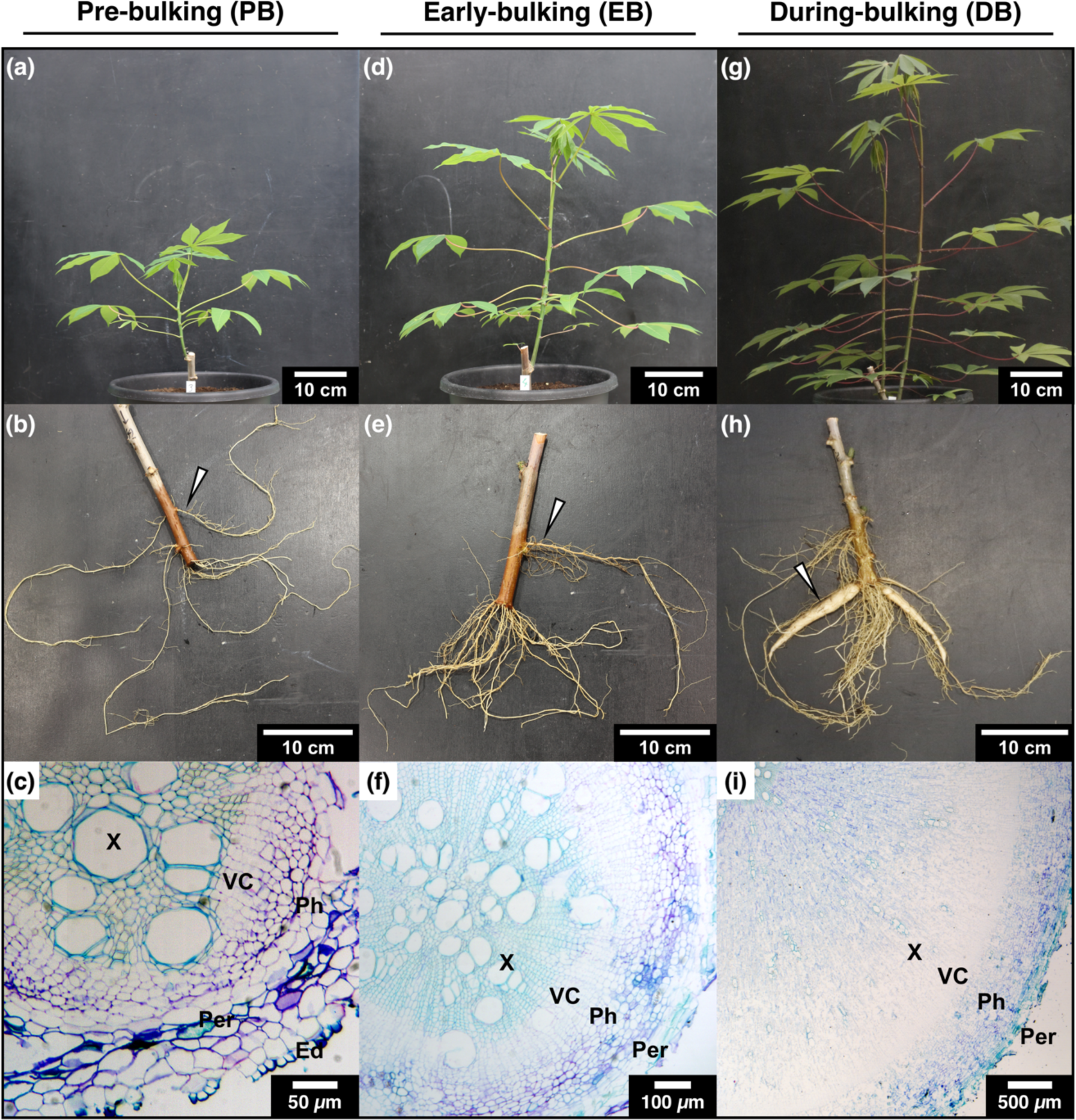
Early development of glasshouse-grown cassava. (a), (b), (c) Pre-bulking ∼30-38 dap. (d), (e), (f) Early-bulking ∼42-51 dap. (g), (h), (i) During bulking ∼60 dap. (a), (d), (g) Shoots. (b), (e), (h) Root stocks. (c), (f), (i) Micrographs of paraffin embedded storage roots. Arrows in (b), (e), (h) depict the thickest nodal root, which was used for determination of the developmental stage. Abbreviations: Ed: Epidermis, Per: periderm, Ph: phloem, Pi: pith, VC: vascular cambium, X: xylem

Under the growth conditions used for this study, the PB stage is usually reached between 30 to 38 days after planting (dap). At this stage the plants are small (Fig. 1a) and have no visually distinguishable storage root developed (Fig. 1b, c). At around 42-51 dap the first clearly thickened roots were visible (Fig. 1e). These were of darker color due to the newly formed periderm (Fig. 1f). At around 60 dap the plants had grown considerably (Fig. 1g), showed the typical red colored petioles of the genotype TME7, and developed clearly thickening storage roots (Fig. 1h, i). PB storage roots did not show a fully annular vascular cambium and the first periderm cells were visible under the still intact epidermis or completely missing (Fig. 1c). Fitting their darker color, the epidermis of EB roots was fully replaced by the periderm (Fig. 1f). The vascular cambium was circular in shape but did not produce the characteristic storage parenchyma yet. DB roots were much larger in diameter than roots of the previous stages (Fig. 1h). Most of this expansion was due to the production of xylem storage parenchyma cells by the rapidly dividing vascular cambium (Fig. 1i). Numerous xylem rays – every second to fourth cell layer – were already established at PB (Fig. 1c). During PB and EB roots did produce many xylem fibers and complex secondary tracheary elements (Fig. 1c, f). The final storage root did only produce very few such lignified cells and mostly simple tracheary elements (Fig. 1i).

Adult stems and storage roots were engulfed in a smooth periderm and were of round shape, mainly differing in the distribution of woody tissue in the xylem and phloem area. The storage root did not produce many tracheary elements and almost no fibers (Fig. 2c), while the lower stem displayed the opposite phenotype (Fig. 2b). Using iodine staining, starch could be detected in the storage root in almost every cell outside the vascular cambium (Fig. 2f). The stem showed iodine staining both in the xylem rays and the axial parenchyma cells connecting them (Fig. 2d,e). Phloem/cortex parenchyma in the stem were green before embedding, indicating the presence of chloroplasts. The storage root cortex contained amyloplasts rather than chloroplasts (Fig. 2f). Interestingly, the pith tissue of the adult stem was also parenchymatic and contained many starch granules (Fig. 2e), thereby resembling the storage parenchyma of the storage root. This pith storage parenchyma was only found in the oldest stem parts, whereas younger stem regions showed larger cells with small secondary cell walls and no starch granules (Fig. 2a, d). This pith tissue made up a larger part of the total stem area, had a foamy consistency, and was hydrophobic. At PB most of the lower stem wood was comprised of storage parenchyma cells in the pith tissue, while the ratio of starchy to woody tissue decreased with time (Fig. S1). This opposing behavior of storage root and wood was further visible in the abundance of major NSC (Fig. 3; Supplementary file 1).

**Fig. 2.**
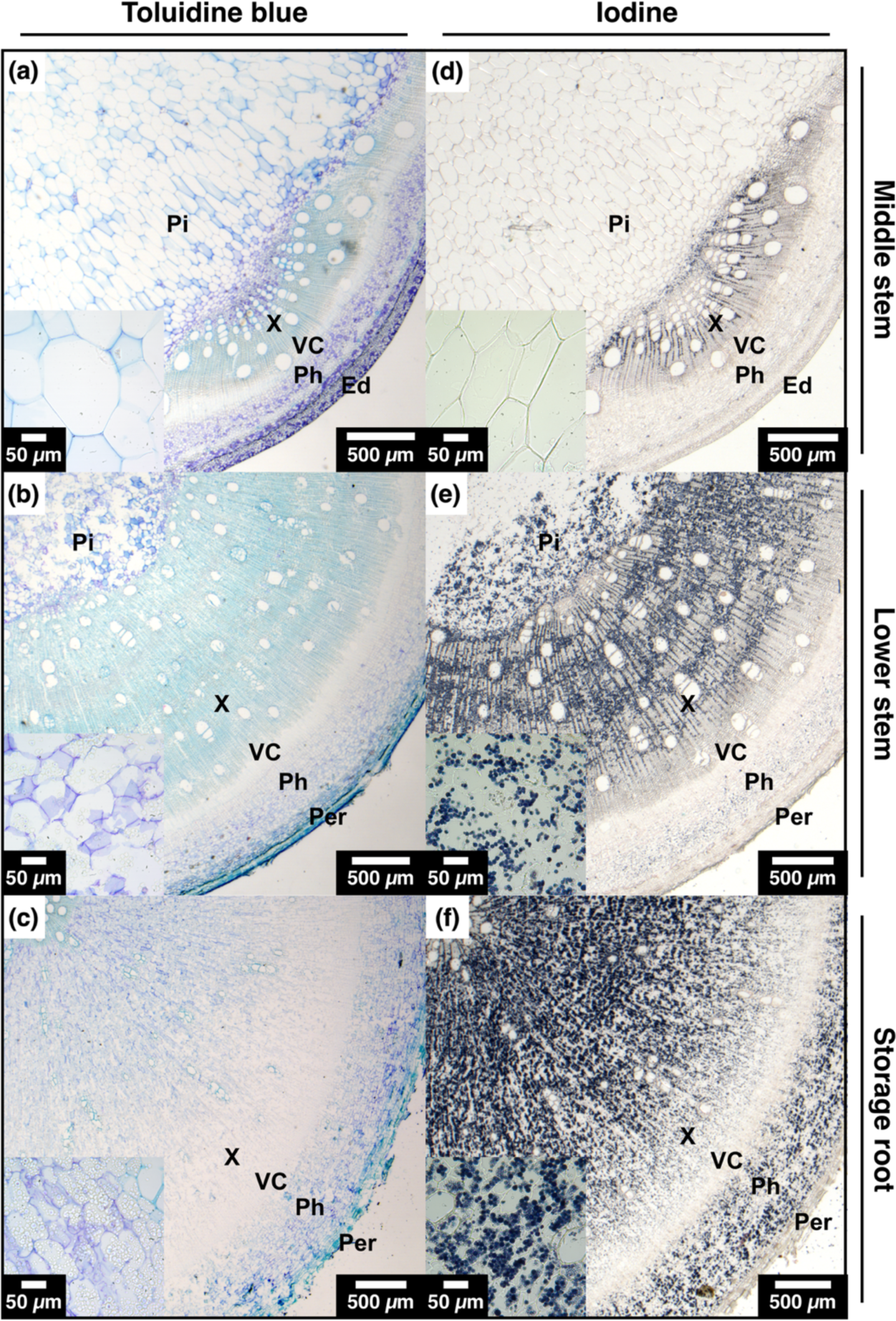
Comparison of cassava stems and storage root during bulking. Stereo micrograph of cross sections stained with either toluidine blue (left column) or iodine (right column) from paraffin embedded green middle stem (a), (d), brown lower stem (b), (e), or storage root (c), (f) samples. Inlays show a micrograph of pith parenchyma of the stem (a), (b), (d), (e), or xylem storage parenchyma of the storage root (c), (f). Abbreviations: Ed: Epidermis, Per: periderm, Ph: phloem, Pi: pith, VC: vascular cambium, X: xylem.

**Fig. 3.**
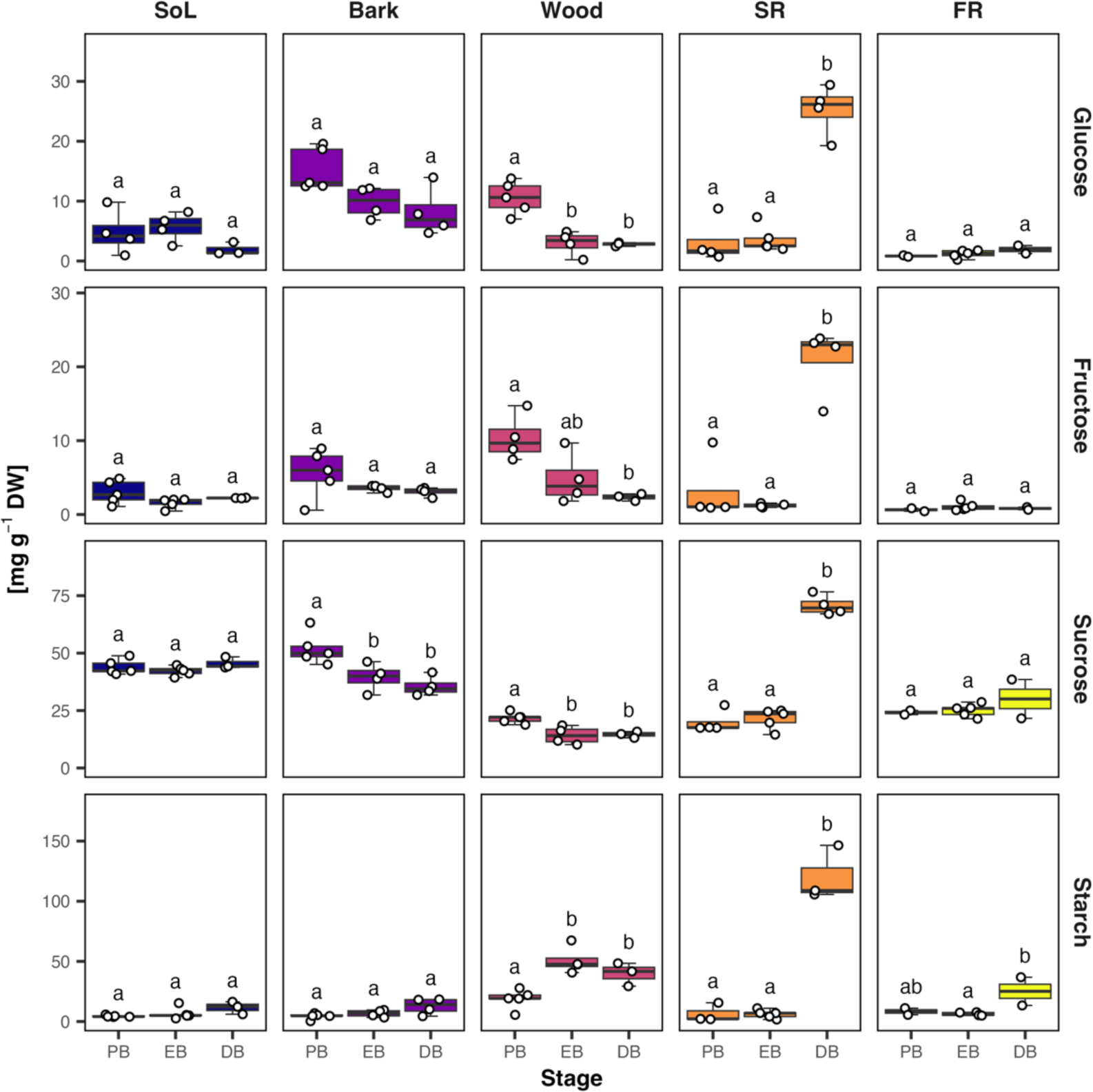
Abundance of major NSC in different casava tissues across developmental stages. Rows from top to bottom show glucose, fructose, sucrose, and starch content in milligram per gram of dry weight. Starch content is shown as the amount of glucose units after amylase/amyloglucosidase digestion. Columns and colors indicate different tissues. A generalized linear model was fitted on the raw values versus the stage for each tissue. Letters show groups of significance within each facet (Tukey HSD p ≤ 0.05) of models with an FDR adjusted p value < 0.05 in an ANOVA. Abbreviations: FR: fibrous root, SoL: source leave, SR: storage root.

Soluble sugars in the storage root rapidly accumulated upon root bulking (DB), while they decreased in the stem wood, indicating a change in carbon allocation behavior. Starch levels strongly increased only at DB in storage root, while they initially increased in the stem xylem and subsequently dropped between EB and DB. The total number of starchy xylem parenchyma cells still increased at this point (Fig. S1). Hence, the absolute starch content should increase over time. In accordance with the iodine staining (Fig. S1), starch content was highest in the wood before root bulking. In bark tissue, only a slight decrease was measured for the soluble sugars, which was significant only in the case of sucrose (Fig. 3). No changes in the soluble sugar contents of the leaves between the different timepoints were apparent. In both leaf and bark tissues, starch increased at DB, but this was not statistically significant (Fig. 3). NSC levels of the fibrous root was largely invariant, but a small increase in starch content was also observed at DB.

### Generating a source-sink transcriptome and phylogenies of selected gene families

Five cassava plants were harvested at three different timepoints. The developmental stage of each harvested plant was determined by microscopy of the storage root. Three PB (38 dap), five EB (51 dap), and four DB (60 dap) plants could be verified to be in the desired developmental stage. Source leaves, the bark and wood of the oldest stem parts, fibrous roots and storage root were sampled. Isolated total RNA of these samples was sent for mRNA sequencing.

In a dimensional reduction projection of the RNA-seq data (UMAP; Fig. 4a), the samples mainly clustered according to their tissue, but some differences between stages could be observed. In particular, the storage root samples of the DB stage were distant from PB and EB, which highlights the strong changes the root underwent during the bulking process. Differentially expressed genes (DEGs) were clustered according to their expression profiles in each tissue (Fig. 4b, c; Supplementary file 2). The majority of genes displayed significant expression changes in the storage root (12,188), while comparably few genes were differentially expressed in source leaf (3,098), bark (3,008), or wood (1,882) tissue. 8,051 DEGs were observed in fibrous root samples, but only a few showed higher expression in DB compared to the other stages (cluster FR 0). Nevertheless, some genes did overlap with the corresponding storage root cluster (SR 4). These genes were involved in starch metabolism, which was in line with the slight increase in starch levels also observed in fibrous root at DB (Fig. 3). However, the overlap between storage root and fibrous root genes with the same expression profile was small. 7,126 DEGs of the fibrous root were part of FR 1 or 2, which showed a drop or rise in expression at EB (Fig. 4d), something that was not found in storage root. Four out of five storage root clusters (SR 1-4) were roughly the same size with 2,400-3,312 DEGs. SR 2 and 3 included genes that were generally lower expressed in later stages, while SR 1 and 4 displayed the opposite behavior.

**Fig. 4.**
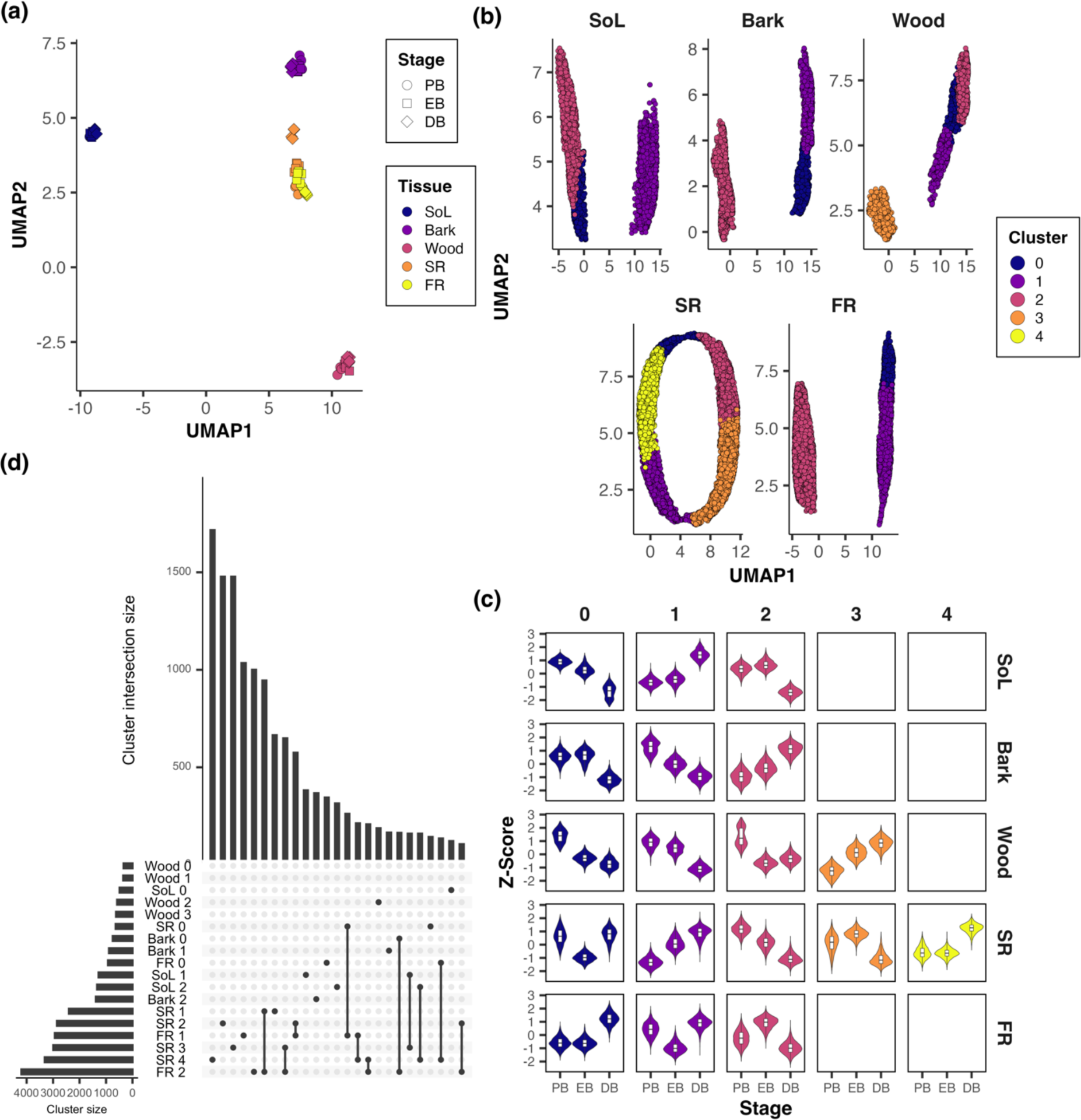
Basic description of the generated cassava transcriptome data. (a) UMAP projection of samples based on all transcripts (DESeq2 normalized counts > 50). (b) Visualization of a CRN analysis for DEGs (FDR < 0.001) within a tissue using a UMAP projection. Each point represents a single gene across all samples. Colors indicate the different clusters found through community detection. (c) Expression profiles of all data points for each gene within a cluster (columns) within each tissue (rows) shown as violin plot. (d) UpSet plot depicting cluster size (horizontal bars) and size of the 25 largest intersections (vertical bars). Abbreviations: FR: fibrous root, SoL: source leave, SR: storage root.

To specifically study the expression of genes involved in phloem loading, carbon transport and unloading, as well as subcellular carbon and water transport, maximum likelihood trees of target gene families (Table 1) from *A. thaliana*, *P. trichocarpa*, and *M. esculenta* were produced (Supplementary files 3-13), cassava genes were annotated accordingly (Supplementary file 14), and the tissue-specific expression of these genes was analyzed.

### Phloem loading and unloading

Phloem loading of sucrose is heterogeneous in nature (Slewinski et al., 2013), but can generally be split into three distinct strategies: (1) Active apoplasmic, (2) passive symplasmic, and (3) active symplasmic/polymer trapping (Rennie and Turgeon, 2009). To this end, phylogenetic trees of *SUGARS WILL EVENTUALLY BE EXPORTED TRANSPORTER* (*SWEET*) (Fig. 5a) and *SUCROSE TRANSPORTER* (*SUC/SUT*) (Fig. 5b) genes were generated and their expression profiled. Active apoplasmic loading requires facilitated export of sucrose from the mesophyll into the apoplast via SWEET11/12 (Chen et al., 2012) and active import via SUT1/SUC2 into the companion cell (CC) (Gottwald et al., 2000). Of the annotated SWEET encoding genes (Fig. 5a), no expression of the *AtSWEET11-14* orthogroup nor any other sucrose transporting clade III *SWEET* was found in the leaves of cassava (Fig. 5c). Furthermore, the expression profile of the *SUT1* orthologs more closely resembles passive symplasmic tree species compared to apoplasmic loaders. For example, in the active apoplasmic loader *A. thaliana, AtSUC2/SUT1* is specifically expressed in CC (Truernit and Sauer, 1995) and is commonly used to facilitate the CC specific expression of transgenes, while orthologs in tree species are mainly expressed in the wood in addition to the phloem (Decourteix et al., 2008, Dobbelstein et al., 2019). Expression of both *MeSUT1* orthologs decreased over time in the storage root and bark but increased in the wood (Fig. 5d). *MeSUT1b* had much higher expression than *MeSUT1a* and was most abundant in woody tissues. *MeSUT1a* had its highest expression in fibrous root and young storage root. However, *MeSUT1b* expression was still present in the source leaf.

**Fig. 5.**
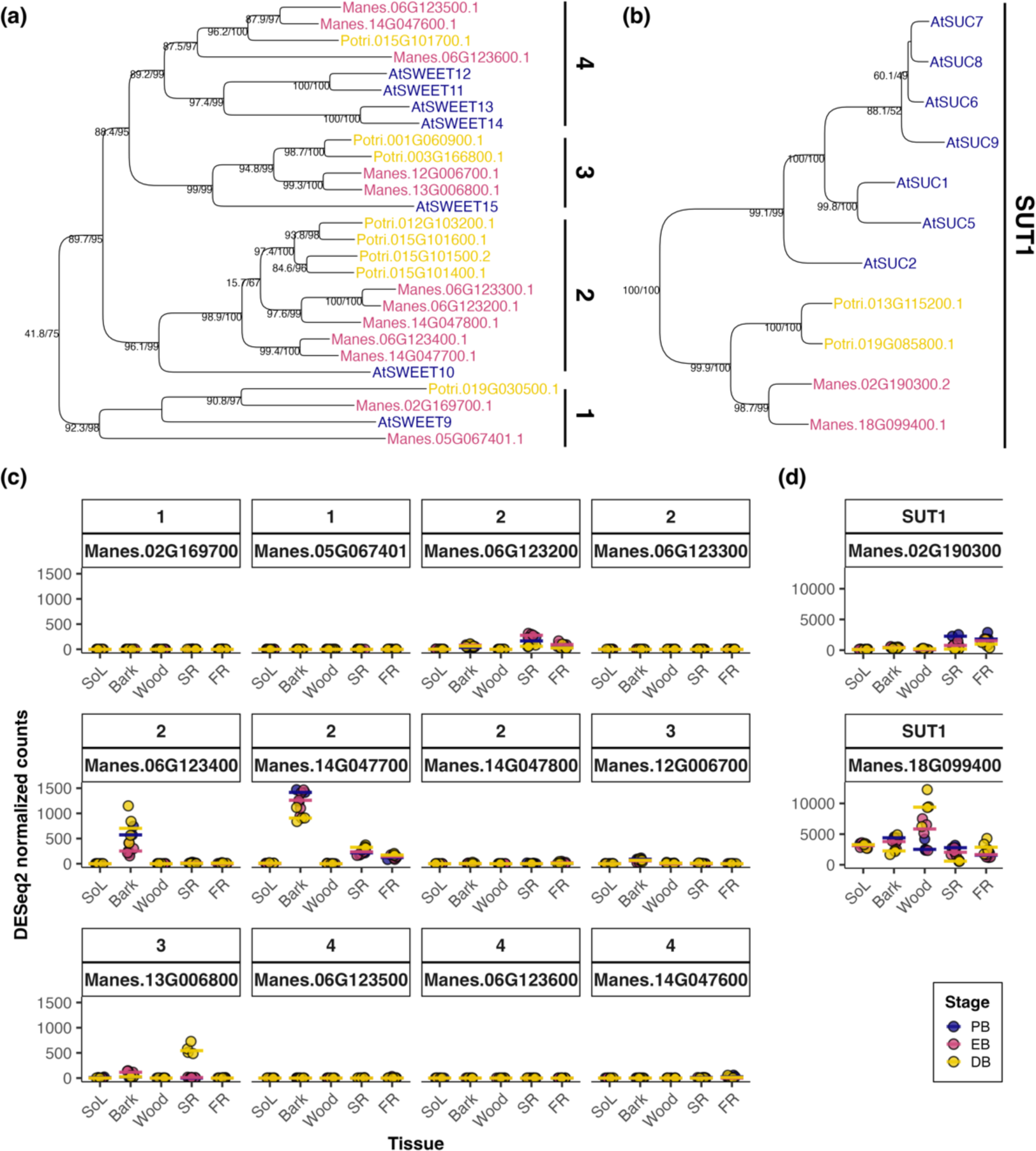
Annotation and expression profiling of cassava SWEET III and SUT1 encoding genes. (a, b) Subtrees of (a) sucrose transporting clade III SWEET and (b) SUT1 encoding genes in *A. thaliana*, *P. trichocarpa*, and *M. esculenta*. Subtrees were taken from midpoint-rooted maximum likelihood trees of full-length CDS sequences (see supplementary data). Branch labels indicate ultra-fast bootstrap support in per-cent and shlrt results. Trees were generated using IQTree. (c, d) Scatterplot displaying the expression levels of cassava (c) *SWEET* clade III and (d) *SUT1* orthologs. Colors indicate the developmental stage. Horizontal lines show the median per developmental stage. Legend is shared between (c) and (d). Abbreviations: FR: fibrous root, SoL: source leave, SR: storage root.

To gather more information about the loading type of cassava, serial block face scanning electron microscopy (SBFSEM) of cassava minor veins was performed to analyze the presence and anatomy of plasmodesmata in the CC/bundle sheath (BS) interface (Fig. 6a-c). Several symmetrically branched plasmodesmata were present at the CC/BS interface (Fig. 6a, c), characteristic for symplasmic but not apoplasmic loaders (Haritatos et al., 2000, Rennie and Turgeon, 2009), while the CC/SE interface showed the generally typical asymmetrically branched pore plasmodesmata (Fig. 6b). However, plasmodesmata in cassava minor veins were not found throughout the whole CC/BS interface but were arranged in pith fields made visible through SBFSEM (Figure 6d-f; Movie S1).

**Fig. 6.**
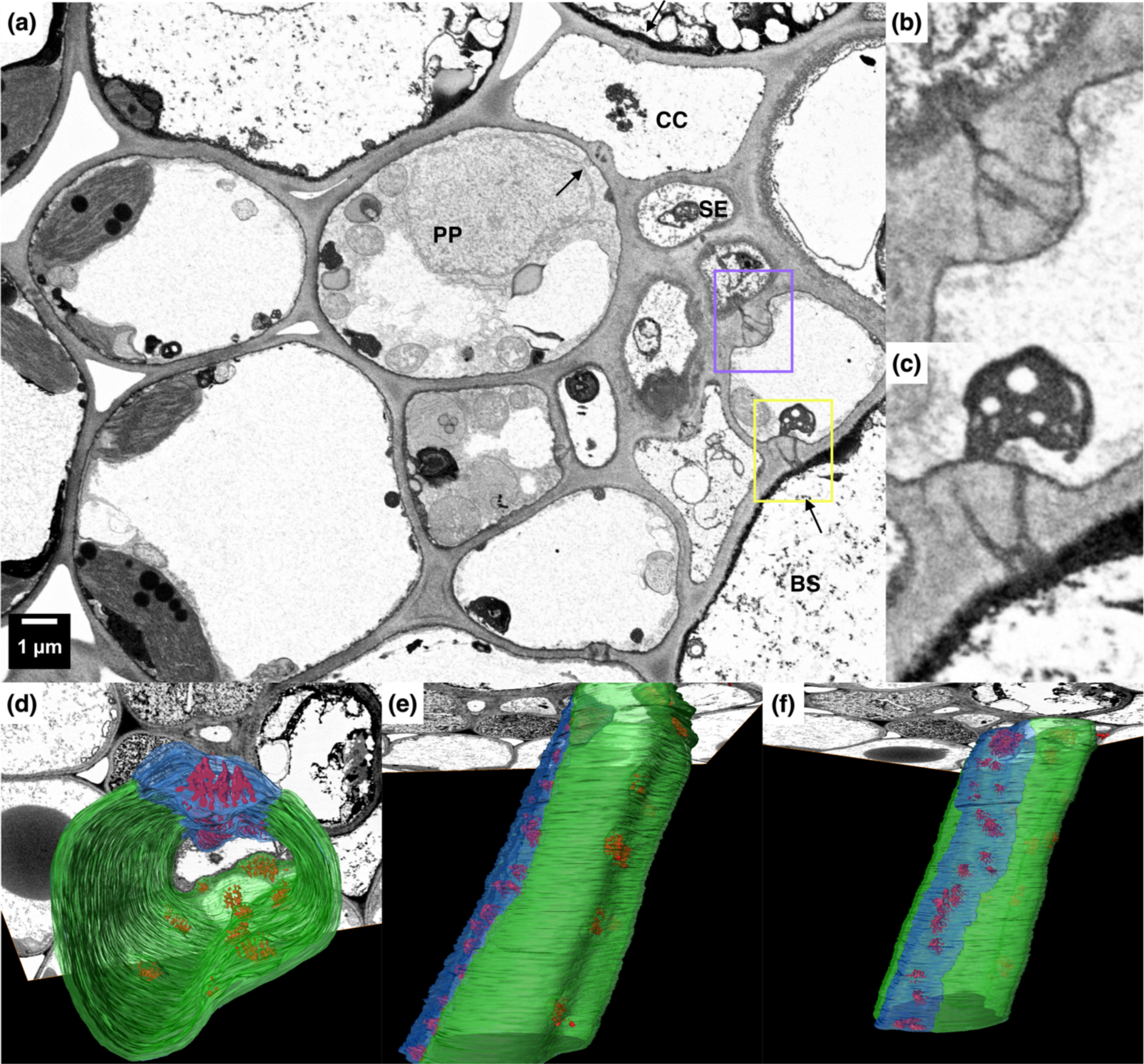
2D and 3D scanning transmission electron microscopy of a cassava minor vein. 2D TEM image of a cassava minor vein. Yellow and purple highlighted areas show a pith field between a bundle sheath and companion cell (b), or sieve element and companion cell (c), respectively. Arrows indicate pith fields between bundle sheath/phloem parenchyma and companion cells. (d-e) 3D reconstruction of a minor vein companion cell. Colors indicate different cell interfaces. Blue: companion cell/sieve element, green: companion cell/bundle sheath. BS, bundle sheath; CC, companion cell; PP, phloem parenchyma; SE, sieve element.

In addition, the sucrose concentration in cassava source leaves is very high with more than 200 mmol l^-1^ at the DB stage (Fig. S3; Supplementary file 16), which is another characteristic of passive symplasmic loaders (Rennie and Turgeon, 2009). An active polymer trapping mechanism is unlikely, due to the absence of specialized intermediary cells, as well as the lack of raffinose in leaf phloem exudates (Fig. S2; Supplementary file 15) (McCaskill and Turgeon, 2007, Rennie and Turgeon, 2009).

Previous studies already showed that the storage parenchyma cell of the storage root and xylem of the stem are symplasmically connected to the phloem via xylem rays (Mehdi et al., 2019), similar to those found in other woody perennials (Van Bel, 1990, Furze et al., 2018). Together with the findings presented here, we conclude that the secondary vasculature and photosynthetically active source leaf cells of cassava are all symplasmically connected (for a model see Fig. 7). In addition to the expression changes of *SUT1* genes (Fig. 5), *SPS* and *SUS* genes (Fig. 7) also showed clear tissue-specific expression changes among the selected and annotated genes (Table1).

**Figure 7.**
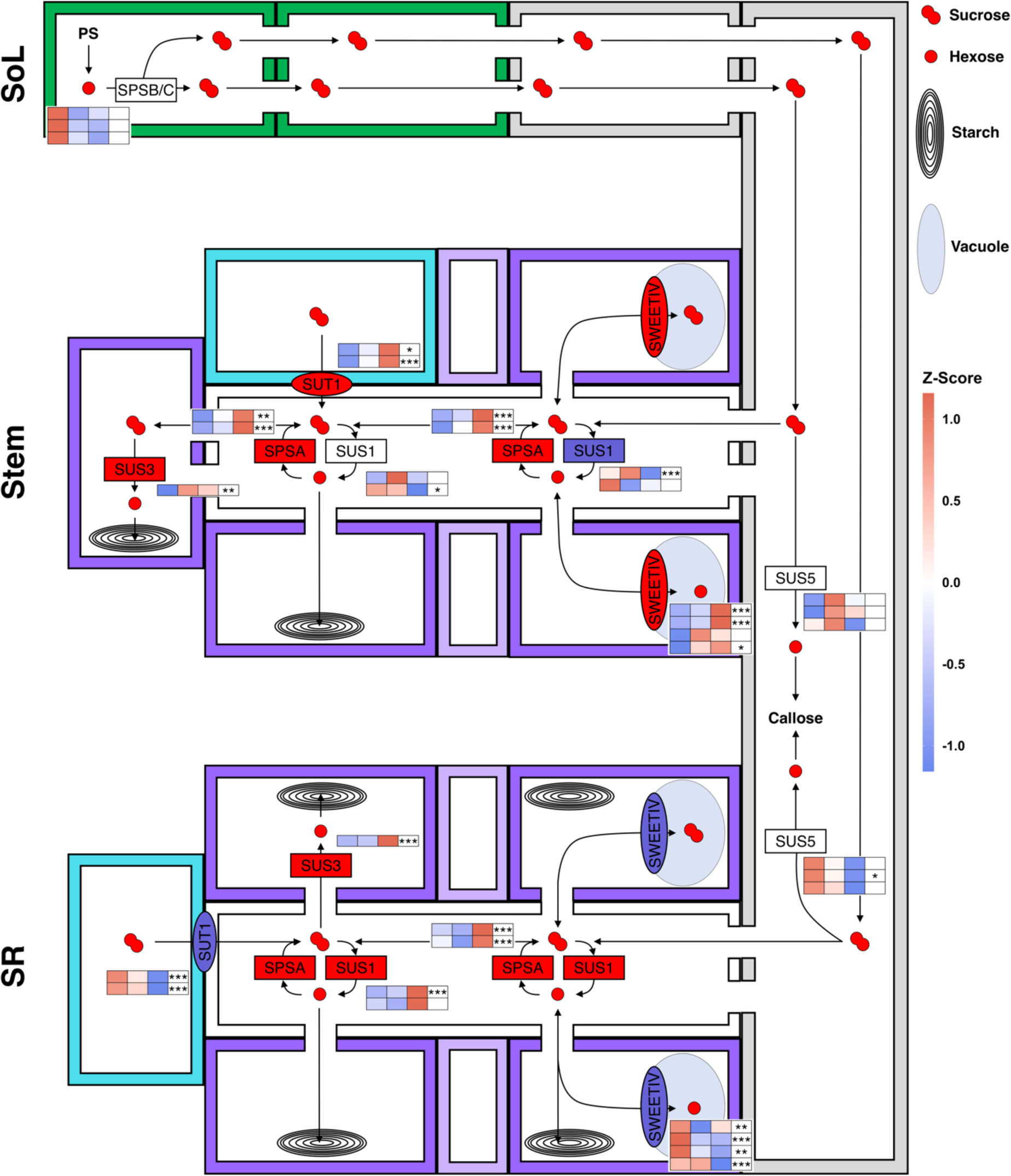
Model of symplastic sugar transport routes in cassava. The top, middle and bottom row depict the source leave, stem, and storage root of the cassava plant. Colored rectangles display idealized cells: grey, companion cell/sieve element complex; white, xylem ray; dark purple, parenchyma cell; light purple, vascular cambium; turquoise, tracheary element. Light blue ellipses are vacuoles, concentric ellipses amyloplasts. Smaller rectangles and ellipses indicate enzymes and transport proteins. Proteins which genes show a general upward or downward trend overtime are colored red or blue, respectively. The heatmaps within the cells display the z-score of the average log transformed values per stage within the given tissue. Stem bark and wood is separated by the vascular cambium. Expression in the storage root is for both xylem and phloem tissue. A likelihood-ratio test was performed with the R package DESeq2 using the intercept only as reduced model. Asterisks indicate significant changes (FDR adjusted p-values; * < 0.05; ** < 0.01; *** < 0.001). Only genes with an average of 50 or higher DESeq2 normalized counts per condition were plotted. Sol, source leave; SR, storage root.

SUS1/4 proteins have previously been shown to be associated with symplasmic unloading (Zrenner et al., 1995, Bieniawska et al., 2007, Gerber et al., 2014, Mehdi et al., 2019, Yao et al., 2019). Of the *MeSUS1* genes that could be annotated, *MeSUS1a* had the highest transcript levels and was differentially expressed in bark and storage root. It was grouped into cluster Bark 0 and SR 4 (Fig. 4, Supplementary File 2), meaning transcript abundance decreased at DB in bark, but increased in storage root (Fig. 7). Overall, *MeSUS1a* was most highly expressed in the wood and adult storage root. *MeSUS1b* showed the same tendencies, but did not reach statistical significance, except in source leaf samples, where it decreased at DB (cluster SoL 0). The temporal expression profile of the *MeSUS1* orthologs did not match starch levels, implying a more general role of this isozyme in ensuring carbon availability for the sink tissues as described in poplar (Coleman et al., 2009, Gerber et al., 2014). However, *MeSUS3* did follow starch content more closely within each tissue. In *A. thaliana,* SUS3 is associated with seed endosperm tissue, which could imply a role for storage (Yao et al., 2019). In addition to *MeSUS1*, *MeSPSA* genes increased in transcript abundance in bark and storage root (Fig. 7), but also in the other tissues (Supplementary File 14). Sucrose cycling in sink organs is a commonly observed phenomenon, potentially explaining the changes in SPSA expression (Geigenberger and Stitt, 1993, Nguyen-Quoc and Foyer, 2001). The other *MeSPSA* gene was similarly expressed across tissues, while transcripts for genes encoding for *MeSPSB/C* proteins were much more abundant in source leaf, suggesting their involvement in photosynthetic sucrose production (Fig. 7). Another interesting aspect is the tissue-specific regulation of genes encoding for tonoplast localized clade IV SWEET proteins (Klemens et al., 2013, Guo et al., 2014). SWEETIV proteins are involved in wood formation in *A. thaliana* (Aubry et al., 2021). In keeping with the Arabidopsis data, increased expression was observed in cassava bark tissue upon increased wood formation at DB and a decrease in *MeSWEETIV* expression was observed in storage root when barely any wood is formed. Generally, changes in the expression of tonoplast sugar transporters were among the more obvious changes in the sugar-related transcripts of the DEGs found in the storage root.

### Compartmentation of metabolites in the bulking storage root

Most known genes involved in vacuolar sugar import were more highly expressed in later stages or at DB. These include all four *TONOPLAST SUGAR TRANSPORTER* (*MeTST*)*1/2* that were all part of cluster SR 4. The only expressed *TST3* orthologue (*MeTST3b*) increased throughout bulking (cluster SR 1). TSTs are H^+^/hexose antiporters (Wormit et al., 2006, Wingenter et al., 2010) but TST1 and TST2 can also facilitate the transport of sucrose into the vacuole (Schulz et al., 2011, Jung et al., 2015).

Other genes grouped in clusters SR 1 and 4 include four out of five *VACUOLAR PYROPHOSPHATASE* (*MeAVP*)*1*, as well as 26 of 35 genes encoding for MeV-ATPase subunits. Both of these proteins allow for energization of the vacuole (Dietz et al., 2001, Gaxiola et al., 2001). The remaining V-ATPase orthologs were either not clustered or not expressed in the storage root.

Conversely, transcripts of proteins involved in vacuolar export such as the H^+^/sucrose symporter MeSUT4b (Schneider et al., 2012), as well as all three *EARLY RESPONSE TO DEHYDRATION 6* (*MeERD6*) proteins (Kiyosue et al., 1998), and two out of three *MeEDL4/ERDL6* orthologs (Poschet et al., 2011) decreased throughout bulking (cluster SR 2 or 3). ERD6 and ERD6-like proteins are part of the MST family and facilitate H^+^/hexose symport across the tonoplast (Pommerrenig et al., 2018, Khan et al., 2023).

In addition to NSC content (Fig. 3), metabolite profiles of EB and DB samples were measured. These showed an increase in *myo*-inositol and other putative vacuolar metabolites (Supplementary File 17). INT1 is the only known protein to be able to export *myo*-inositol from the vacuole (Schneider et al., 2008). The sugar alcohol - *myo*-inositol, and indeed all soluble NSCs, is/are used by plants to increase the osmotic potential of the vacuole. Accordingly, changes in the expression of genes relevant for water transport were also observed. Passive transport of water molecules across membranes is enabled by AQUAPORIN (AQP) family proteins. Plasma membrane and tonoplast localized AQPs are called PLASMA MEMBRANE INTRINSIC PROTEINS (PIPs) and TONOPLAST INTRINSIC PROTEINS (TIPs), respectively. In poplar another plasma membrane localized group of AQPs was identified (XIPs; Lopez et al. (2012), Maurel et al. (2015)), but, while present, no members of this subfamily were expressed in cassava. However, nine of 13 expressed PIPs and five of nine expressed TIPs were significantly increased in expression throughout root bulking (cluster SR 1 or 4).

Due to the high starch content of storage parenchyma cells, expression of genes involved in carbon allocation into amyloplasts was also analyzed. Interestingly, all four *GLUCOSE-6-PHOSPHATE/PHOSPHATE TRANSLOCATOR* (*GPT*) genes sharply increased in their abundance at DB in storage root (cluster SR 4, Fig. 8). GPT supplies the amyloplast with glucose-6-phosphate (G6P) in exchange for phosphate (Kammerer et al., 1998). The G6P isomer G1P is a precursor to ADP-glucose that is the substrate for starch biosynthesis. During the conversion of ADP-glucose to starch ADP is produced which is exchanged with ATP by a NUCLEOTIDE TRANSPORTER (NTT) (Reiser et al., 2004). Two orthologs of *NTT* could be annotated in cassava, one was co-expressed with the *GPT* orthologs (Fig. 8). The expression of the only annotated plastid sucrose transporter (*PLASTID SUGAR TRANSPORTER, pSUT*) was decreased upon root bulking, while one of two *PLASTID GLUCOSE TRANSPORTERS* (*pGlcT*) showed the opposite behavior. Both of these transporter types export their respective substrates from the plastid to the cytosol (Cho et al., 2011, Patzke et al., 2018). While it is not fully clear, where the H^+^/glucose antiporters *SUPPRESSOR OF G PROTEIN BETA1* (*SGB1*) are localized within the cell, they share a phylogenetic group with *pGlcT*. Three out of four *MeSGB1* orthologs were found in cluster SR 1, as they were higher expressed throughout root bulking.

**Fig. 8.**
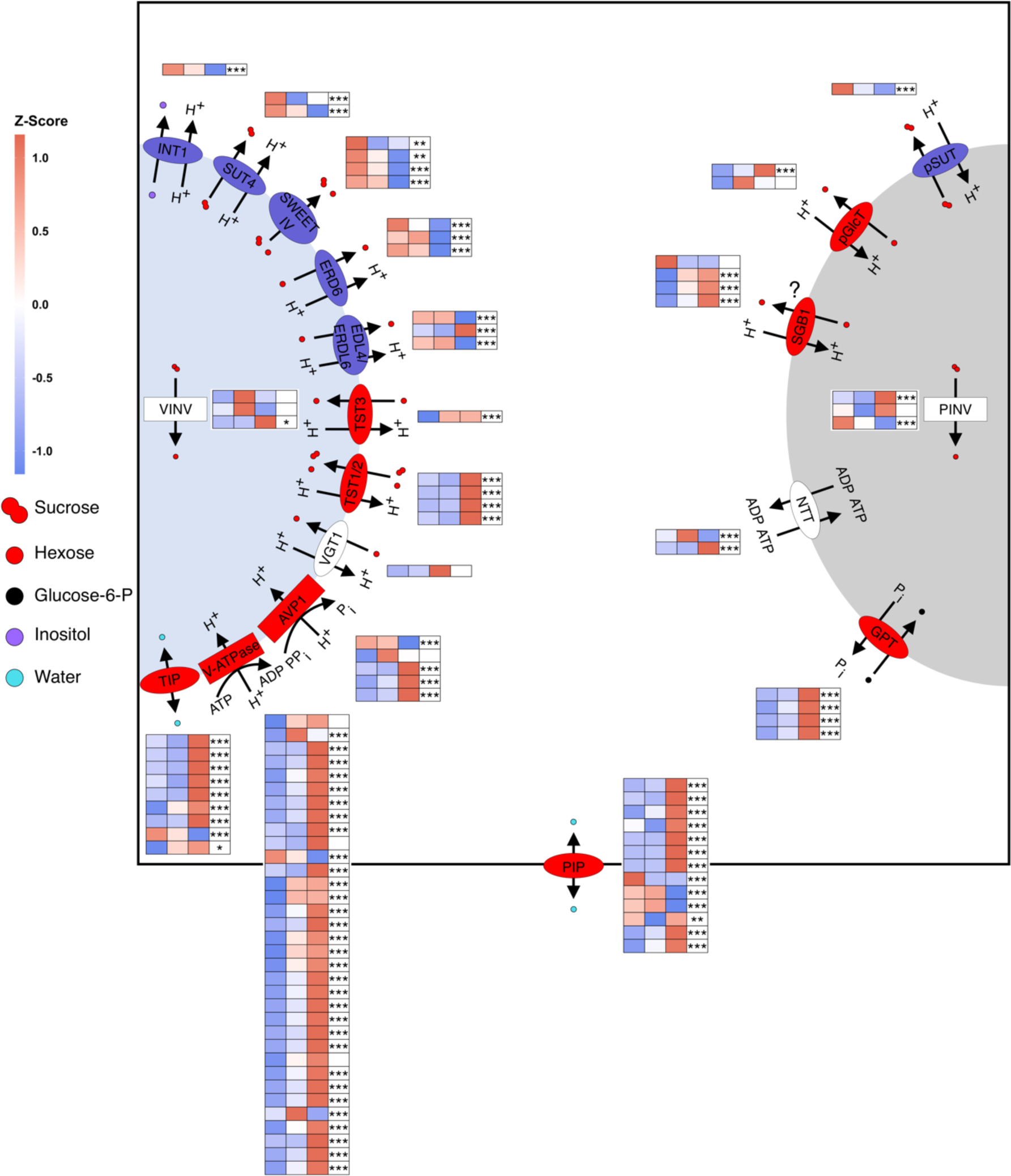
Vacuolar and amyloplast transporters in cassava storage roots throughout bulking. The blue and grey half-circle represent the vacuole and amyloplast, respectively. The black border shows the plasma membrane of a parenchyma cell. Rectangles and ellipses indicate enzymes and transport proteins, respectively. Proteins which genes show a general upward or downward trend overtime are colored red or blue, respectively. Heatmaps show the z-score of the average VST value per stage in the storage root of two independent experiments. Only genes with > 50 normalized counts on average are shown. From left to right: Pre-bulking, ∼30-38 dap; early-bulking ∼42-51 dap and during bulking ∼60+ dap. A likelihood-ratio test was performed with the R package DESeq2 using the intercept only as reduced model. Asterisks indicate significant changes (FDR adjusted p-values; * < 0.05; ** < 0.01; *** < 0.001).

To directly measure metabolite compartmentation within the storage root, non-aqueous fractionation (NAF) was performed to compare NSC content of the cytosol, vacuoles, and plastids (Fig. 9). Unfortunately, only adult storage roots could be measured successfully. In these, glucose, fructose, and sucrose showed the same relative distributions across the different compartments (Fig. 9), but sucrose was much more abundant (Supplementary file 18). As suggested by the changes in gene expression of vacuolar transporters (Fig. 8), NSC abundance was highest in the vacuole with approximately 40% of the total content, however plastids were a close second with around 35%. As expected, most of the sugar accumulating in the storage root was not present in the cytosol but was compartmentalized into other subcellular compartments. The presence of sucrose in amyloplasts was additionally described for potato tubers, however, whilst a chloroplast transporter capable of transporting sucrose has been identified (Patzke et al., 2018), it is as yet unclear how sucrose enters amyloplasts (Hampp and Schmidt, 1976, Gerrits et al., 2001). The presence of hexoses in the plastid can be explained through plastid A/N-INV proteins (Gerrits et al., 2001). These genes were all expressed, but no DEGs were found between the different tissues and timepoints, respectively (Fig. 8). Overall, the data indicates pronounced regulation of subcellular transport during storage root bulking.

**Fig. 9.**
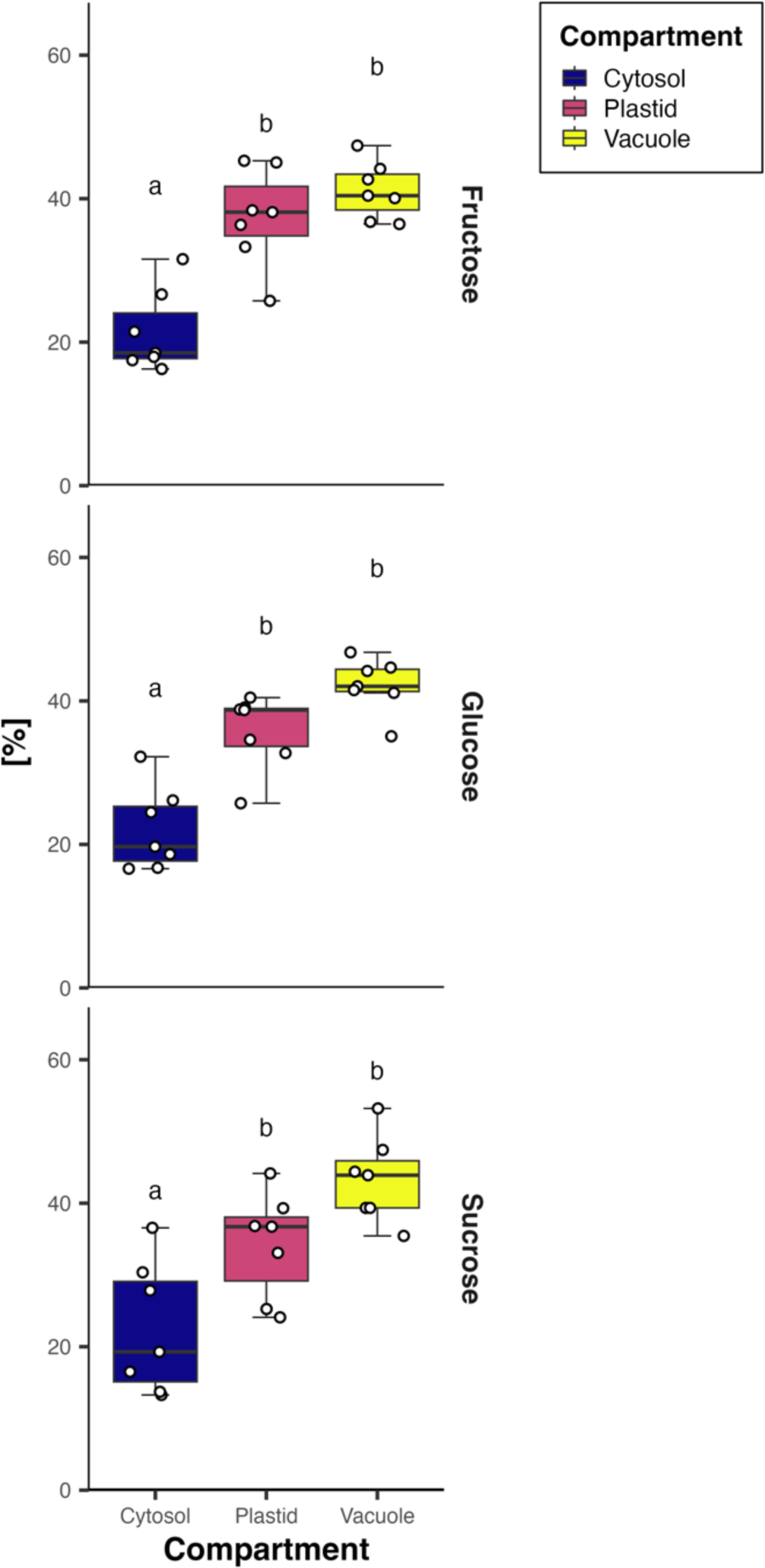
Non-aqueous fractionation and sugar measurement of adult cassava storage roots. Data is displayed as per cent of total sugar content. Samples are independent from the other experiments. Rows from top to bottom show fructose, glucose and sucrose content. A linear model was fitted using predicting the relative sugar quantities per compartment across 7 replicates. Letters show groups of significance within each facet (Tukey HSD p ≤ 0.05) of models with an FDR adjusted p value < 0.05 in an ANOVA.

## Discussion

### Cassava stems are large carbon sinks with complex development

Stems of any type of plant are no mere highways for metabolite and nutrient transfer, but are rather important and complex carbon sinks, especially in woody perennials such as cassava (Furze et al., 2018). Even in the presented short time glasshouse study, the plant stems contained more than five percent of their total DM as starch, while also producing the necessary secondary cell walls (Fig. 3). The tissue composition of cassava stems also appeared to be connected to the initiation of the storage root (Fig. 2, S2). The first newly formed stems were largely made up from large pith parenchyma cells, which were filled with starch. These cells apparently reach their full capacity already during the EB stage. Coincidentally, this was around the same time the first starch granules could be found in xylem rays and cortex parenchyma of developing storage roots (Fig. S2). Hence, induction of the storage root could function as an “overflow basin” for unmetabolized sugar of the stem, which would explain why root bulking initially occurs proximally to the shoot. It is likely that sugar signaling plays an important role in storage root initiation as starch metabolism in the root occurs earlier than storage parenchyma formation. Stem starch metabolism, however, is not limited to the pith parenchyma, but can be observed in xylem rays as well as axial parenchyma, which is the usual xylem parenchyma pattern in different woody species (Sokołowska and Zagórska-Marek, 2012, Furze et al., 2018).

### Parenchymatic cells of source and sink tissues are symplasmically connected

Phloem loading can be grouped into three categories. Apoplasmic phloem loading is commonly known from herbaceous species such as *A. thaliana* but also root and tuber crops such as potato, where sucrose is transported from the phloem parenchyma/transfer cells into the surrounding cell wall space by clade III SWEETs (Chen et al., 2010, Chen et al., 2012) and subsequently taken up into phloem companion cells by SUT1 proteins. However, most woody plants primarily use a passive symplasmic mode of phloem loading (Rennie and Turgeon, 2009, Slewinski et al., 2013, Zhang et al., 2014). Expression of phloem loading *SWEET11/12* can typically only be measured in the leaves and the phloem of apoplasmic loaders (Chen et al., 2012, Abelenda et al., 2019, Doidy et al., 2019), but not in passive loaders (Zhang et al., 2020). Three *SWEET* genes were found in the same orthogroup (*SWEET-III-4*) as *AtSWEET11/12* (Fig. 5a, Supplementary file 14), but none showed expression in any tissue including the source leaf, nor was any other putative sucrose transporting clade III *SWEET* expressed in the source leaf (Fig. 5c). These findings suggest a predominantly symplasmic loading strategy. Additionally, *MeSUT1* expression profiles more closely resembled the pattern of passive symplasmic loading woody perennials, which utilize SUT1 proteins for sucrose uptake from xylem vessels during remobilization, rather than for phloem loading (Loescher et al., 1990, Decourteix et al., 2006). These genes are most highly expressed in bark and wood tissue (Decourteix et al., 2008, Decourteix et al., 2006, Dobbelstein et al., 2019) and their mRNAs are detectable in xylem parenchyma (Decourteix et al., 2008), but expression is still high in phloem tissue (Schrader et al., 2004). *MeSUT1b,* specifically, was more highly expressed in bark and wood tissue, as well as in the source leaf. However, it showed an opposing expression pattern in wood and storage root coinciding with the ratio of woody xylem to storage parenchyma (Fig. 7), implying an expression in vessel associated or regular xylem parenchyma cells like in other species and not in the storage parenchyma (Decourteix et al., 2008).

Symplasmic and apoplasmic loaders differ in the number and structure of their plasmodesmata of the CC/BS or CC/PP interface (Turgeon and Medville, 2004). Many symmetrically branched plasmodesmata could be observed in CC/BS interface of cassava minor veins in SBFSEM images, which is a characteristic of symplasmic loaders, whereas apoplasmic loaders have no or few asymmetrically branched plasmodesmata. Cassava, however, showed an apparently lower frequency than many tree species (Fig. 6c,e) as plasmodesmata were only abundant in very few two dimensional TEM images. SBFSEM that allows for continuous imaging of sequential sections through an embedded specimen to generate 3D volumes, however, revealed the presence of highly branched plasmodesmata at this interface (Fig 6f-h, Movie S1). This highlights the importance of 3D reconstructions when evaluating loading types by anatomical features. Transfer cells, a specialized PP cell type with wall ingrowths towards the CC (Haritatos et al., 2000) that can occur in apoplasmic loaders, were also not found in cassava minor veins, neither were intermediary cells that exist in active symplasmic loaders. Furthermore, sucrose was far more abundant in phloem exudates compared to raffinose (Fig. S2). Notably, measured source leaf sucrose concentrations exceed 200 mmol l^-1^, which is comparable to other known passive loaders (Rennie and Turgeon, 2009, Slewinski et al., 2013). In conclusion, the minor vein anatomy, presence of symmetrically branched plasmodesmata in the CC-BS interface, high leaf sucrose concentration in the source leaves, as well as the lack of expression of the necessary transporters, makes it likely that cassava predominately utilizes a passive symplasmic phloem loading strategy when transferring carbon from leaf mesophyll cells to sink organs.

However, active sucrose uptake into CCs is still plausible due to expression of *MeSUT1* orthologs also in phloem-containing tissues in cassava. Yet, without the expression of sucrose-transporting SWEET proteins, it is unclear how much contribution this active mechanism can have to the overall source-sink carbon allocation. It is, however, still possible that active phloem loading serves an important purpose for apoplastic sugar retrieval under specific conditions that are not apparent under our greenhouse study conditions.

Symplasmic unloading strategies are more widespread among different types of plants, varying even between and within a tissue depending on its development (Viola et al., 2001). Arabidopsis roots, potato tubers, as well as the secondary vasculature of woody perennials, including cassava stems and storage root unload symplasmically (Sokołowska and Zagórska-Marek, 2012, Ross-Elliott et al., 2017, Mehdi et al., 2019). Several studies have shown the involvement of SUS proteins in symplasmic unloading processes. SUS proteins can be grouped into three phylogenetic clades. The SUS1 clade contains the highly expressed *AtSUS1* and *AtSUS4* genes, which encoded proteins that localize to the companion cells (Bieniawska et al., 2007, Yao et al., 2019). However, neither knockout of *AtSUS1* (Barratt et al., 2009), nor knockdown or overexpression of aspen *SUS1* genes alters the macroscopic growth phenotype in glasshouse studies (Coleman et al., 2009, Gerber et al., 2014), likely because symplasmic unloading does not involve CCs at least in *A. thaliana* roots (Ross-Elliott et al., 2017). Beyond Arabidopsis, however, increased SUS activity did lead to increased cellulose content and thickened secondary cell walls in aspen (Coleman et al., 2009), while a decrease reduced the overall carbon availability (Gerber et al., 2014). Furthermore, SUS activity is important for starch production in potato tubers (Geigenberger and Stitt, 1993, Zrenner et al., 1995). Both, the role of SUS in secondary cell wall formation and starch biosynthesis make *MeSUS1* genes interesting research targets. Two *MeSUS1* genes could be annotated (Table S2) that were highly expressed with counts in the high ten-thousands. Their expression was high in secondary vasculature, but strongest in wood and bulking storage root. Previous *MeSUS1a* promoter studies also showed strongest activity in stems and storage roots with notable staining also in transport-associated xylem rays present in both tissues (Zierer et al., 2022). The two *MeSUS1* genes showed an increase in transcript abundance during bulking in storage root but much stronger for *MeSUS1a*, which also decreased at DB in bark, implying a change in sink strength towards the bulking storage root (Fig. 7). In addition to *MeSUS1*, *MeSUS3a* expression was high in the storage root and bark, but also in the wood. Within the wood and the storage root, transcript levels of *MeSUS3a* tightly followed the relative starch content (Fig. 3, 7). This was not true for *MeSUS1a*, implying a role for *MeSUS3a* in starch biosynthesis. *MeSUS5/6* orthologs were only expressed in transport phloem containing tissues (bark, storage root and fibrous root), which fits the observation in *A. thaliana*, where SUS5/6 localize to the sieve elements where they are involved in callose biosynthesis (Barratt et al., 2009, Yao et al., 2019).

In summary, cassava parenchyma cells in source leaf, stem and secondary root tissue seem to form one large symplasmically connected system (Fig. 7). As such, carbon allocation is controlled by the phloem Münch flow and local sink strength might mainly be controlled through growth and carbon storage processes, as well as subcellular carbon compartmentalization.

### Subcellular compartmentation of sugars in the storage root is an important aspect of root bulking

Once the bulking process is initiated, cassava storage roots rapidly increase in diameter. Between the EB and DB stage multiple nodal roots per plant grew from a few millimeters to more than a centimeter in thickness (Fig. 1), while producing over 10% of their total dry biomass as starch (Fig. 3). These changes were accompanied by strong changes in the transcriptome of the storage organ (Fig. 4a), including lowered expression of secondary cell wall producing enzymes, as well as an increase in transcript abundance of starch biosynthesis genes as described previously (Rüscher et al., 2021), but also many proteins involved in compartmentation of sugars and other metabolites (Fig. 8).

The MST gene family includes a plethora of genes encoding for proteins involved in tonoplast or plastid transport (Pommerrenig et al., 2018). While transcripts of proteins involved in vacuolar/plastidic import generally increased when cytosolic sugar levels are high, those involved in vacuolar/plastidic export generally decreased in expression under latter conditions (Wormit et al., 2006, Patzke et al., 2018, Klemens et al., 2013, Khan et al., 2023).

In plants such as sugar beet, orthologs of AtTST2 proteins are involved in sucrose storage in the vacuole (Jung et al., 2015). While cassava storage roots did not reach sugar beet levels of sucrose content, even the small storage roots analyzed in this study reached around 7.5 % sucrose compared to their total DM, which was higher than any other tissue and implies active concentration of the disaccharide. Appropriately, all cassava orthologs of *AtTST1/2* showed the same pattern as sucrose and hexose abundance with a strong increase at DB (Fig. 8). By contrast, homologs of the vacuolar sucrose exporter *AtSUC4* (Schneider et al., 2012) and other vacuolar exporters showed the opposite expression profile (Fig. 8). Direct measurements of sugars through non-aqueous fractionation also suggest that high levels of sugars are accumulated in vacuoles and plastids. Sucrose and the monosaccharides glucose and fructose had the same relative distribution across the compartments and were most abundant in vacuoles, closely followed by plastids, with the cytosol having considerably lower relative sugar levels (Fig. 9). This change in expression of tonoplast transporters was also accompanied by pronounced increase of transcripts encoding for proteins and protein complexes involved in vacuole acidification (especially V-type ATPases; Gaxiola et al. (2007)), likely necessary for maintaining the high levels of secondary active transport across the tonoplast. The apparent increase in cellular- and subcellular osmotic potential during storage root bulking was also associated with an increase in expression of *PIPs* and *TIPs*, which facilitate water inflow into the symplast and vacuoles, respectively.

Taken together, sub-cellular compartmentation appears to be an important aspect of root bulking that has multiple implication on the storage organ’s physiology. For once, the proposed efflux of osmolytes from the symplast in the storage root specifically would create a more favorable pressure gradient towards it, increasing its specific sink strength, hence allowing for more carbon to be allocated to the rapidly growing storage organ (Münch, 1930). Furthermore, the large size of storage parenchyma cells compared to regular xylem parenchyma is probably facilitated through rapid vacuolar swelling comparable to how tracheary elements are formed (Kaiser and Scheuring, 2020). Lastly, the accumulation of osmotic substances, followed by the inflow of water into storage root vacuoles might have evolved as a water storage function to endure the dry season. Regardless, a competition between sugar compartmentalization and its utilization in starch biosynthesis might arise. Direct evidence for this was found recently in Nigerian field trials where a strong negative correlation between sucrose, hexoses and most other measured metabolites with potential osmotic function and dry matter content – itself mainly being attributed to starch concentration – was observed (Gutschker et al., 2023, Lamm et al., 2023), indicating that this trait is not yet fixed in at least this particular cassava population, making it an interesting research and improvement target, even outside conventional breeding.

## Material and methods

### Plant material

Cassava plants (genotype TME7) were planted in 14 l pots from 20 cm long stakes. Five plants were harvested at five timepoints: 25, 30, 38, 51 and 60 dap. The third to fifth source leaf without middle vein, the wooden lower stem, the developing storage roots, as well as an assortment of side- and other fibrous roots were sampled. The lower stem was further split into bark and wood by peeling. storage root samples were defined as a few centimeter-long pieces of the 2-4 thickest, nodal-derived roots, proximal to the stake. The samples were taken in the afternoon around 4pm. For each of the samples a small subsample was put into 30 % ethanol microscopy (toluidine blue and iodine staining). The developmental stage was determined based on the morphology of the thickest part of the furthest developed storage root.

### Paraffin embedding and histology

Specimen were fixed in formaldehyde (3.7 %) / acetic acid (5%) solution in 50 % ethanol (FAA) overnight in a vacuum chamber, infiltrated in a Leica TP20: 1.5 h 70 % ethanol, 2x 1.5 h 90 % ethanol, 1 h 0.5% Eosin-Y in 100 % ethanol, 2x 1.5 h 100 % ethanol, 2x 1.5 h xylol and 2x 24 h paraffin (ParaplastTM Plus, 56 °C melting point) and embedded in a Leica EG1160. A Leica RM2265 was used for sectioning. Paraffin was removed using Roti-HistolTM for 20 min for two changes. The sections were then rehydrated in an ethanol series (100 %, 100 %, 90 %, 80 %, 70 %, 50 %, distilled water) for two minutes each. Staining was performed using toluidine blue (0.1 %) and iodine (5 %).

### Metabolite measurements

Metabolite profiles, as well as sugar and starch measurements, were performed exactly as previously described (Rosado-Souza et al., 2019).

### RNA Sequencing and raw data processing

RNA was extracted using the Spectrum^TM^ Plant Total RNA-Kit (Sigma-Aldrich). DB storage root samples were vortexed for 10 minutes, rather than heated to 65 °C. RNA was sent to a service provider for paired-end mRNA sequencing (> 20 million reads, PE150). Resulting FastQ files were quality controlled using FastQC v0.11.9 (http://www.bioinformatics.babraham.ac.uk/projects/fastqc/) and MultiQC v1.13 (https://multiqc.info/). The reads were trimmed for adapter content and quality via bbduk v38.97) (http://sourceforge.net/projects/bbmap/). Bases under a quality of 30 were trimmed from both sides. Reads shorter than 35 base pairs or with an average quality below 30 after trimming were removed. The trimmed reads were mapped to the cassava genome v8.1. (including plastid and mitochondrial genome) with STAR v2.7.10a (Dobin et al., 2013). Mapped reads were counted using FeatureCounts v2.0.3 (Liao et al., 2014). Only uniquely mapped reads were counted. Multi-overlapping reads were not counted. The expression data for the storage roots was reproduced in an independent RNA-Seq experiment to ensure that these changes really reflect the root bulking process.

### Data analysis

Counts were normalized and log transformed (variance stabilization transformation; VST) using the R package DESeq2 v1.38.3 (Love et al., 2014). The DEG analysis against the developmental stage was performed for each tissue individually utilizing a LRT in DESeq2 using the intercept-only model as comparison. Genes with FDR < 0.001 were accepted as DEGs. For the combined analysis of the two storage root experiments shown in Fig. 8, the VST values were batch corrected using LIMMA v3.54.1 and the LRT executed with the experimental batch included as covariate. Downstream analyses were performed on the scaled and centered VST values. UMAP projection was performed in Python using the umap-learn module v0.5.3. DEGs were clustered through Louvain community detection with resolution 0.75 (Blondel et al., 2008) in python using the NetworkX module v3.0. Edges were drawn between genes based on a Pearson correlation coefficient above a permutation-based threshold (median 99.995 % quantile of 1,000 replications). Genes < 50 edges were removed. For NSC data, generalized linear models were generated based on a Gamma distribution with log link in base R. An ANOVA with type II sum of squares was executed using the car package. All p-values were adjusted for multiple testing using the FDR approach. Relative NAF results were tested using a linear model. Tukey’s HSD was used as post-hoc test for significant results (FDR < 0.05). If not stated otherwise, plots were generated in R using the ggplot2 package and figures prepared in Sketch for macOS.

### Functional annotation

Peptides from the *M. esculenta* genome v8.1 and *P. trichocarpa* genome v4.1 were locally blasted (blast+ v2.13.0) against the *A. thaliana* proteome from the araport11 genome (e < 0.001). A list of aliases was downloaded from TAIR (arabidopsis.org, accessed on April 7^th^, 2021). Protein sequences were scanned against the Pfam database v35.0 using hmmer v3.3.2.

### Phylogenetic analysis

For each protein family, corresponding *A. thaliana* genes were selected via keyword search on arabidopsis.org. Suitable *M. esculenta* and *P. trichocarpa* genes were selected based on BLASTP similarity to *A. thaliana* and presence of a specified Pfam domain (Table 1). Full-length CDS were aligned using the --auto option in MAFFT v7.511 (Katoh et al., 2002) and the alignments trimmed using TrimAl v1.4 (Capella-Gutiérrez et al., 2009). Genes with more than 50% gaps after trimming were removed and the alignment repeated using the l-ins-I algorithm. Maximum likelihood trees were generated using IQ-TREE v1.6.12 (Nguyen et al., 2014) with 1,000 ultra-fast bootstrap repetitions (Hoang et al., 2017) and Shimodaira– Hasegawa approximate-LRT (1,000 repetitions). The substitution model was selected using ModelFinder (Kalyaanamoorthy et al., 2017). Trees were drawn using the R packages ggtree. A full list of annotated genes can be found in the supplementary files (Supplementary file 3-13).

### Serial block face scanning electron microscopy

Leaves harvested from mature *M. esculenta* were cut into 2mm x 2mm pieces and fixed in 4% glutaraldehyde, 2mM CaCl2, 0.1M cacodylate buffer, pH 6.8 at room temperature (RT) for 6 hours. The samples then were fixed in a microwave oven (Biowave Pro, Pelco, Fresno CA, USA) at 350 W at 35°C for 2 minutes. The leaves were washed 3 times 10 minutes with 0.1M cacodylate buffer and then post-fixed overnight at 4°C in 3% potassium ferricyanide, 0.3M cacodylate, pH 6.8, 4mM CaCl_2_, with an equal volume of 4% OsO_4_. The following day the samples were washed 3 times 10 minutes in ddH_2_O at RT, incubated for 40 minutes in 0.5% (w/v) thiocarbohydrazide (TCH) and washed again 3 times 10 minutes in ddH_2_O at RT. The samples were incubated in 2% OsO_4_ at RT and washed 3 times 10 minutes in ddH_2_O after 3 hours. Samples were dehydrated in 10% increments of aceton from 0 – 100% and 3x 100% at RT 10 minutes each step. Following the dehydration, the samples were infiltrated in Hard Spurrs resin without DMAE as follows (1 part resin: 3 parts of acetone overnight, 1:2, 1:1, 2:1, 3:1, 100%, 2x 100% hard Spurrs with DMAE) and microwaved for 1 hour at 100W at 40°C max and cured for 48 hours at 60°C. The blocks were trimmed and mounted as described in (Arcalís et al., 2020). Once the block was mounted on a SEM stub, sections were imaged on an Apreo Volumescope SEM (Thermo Fischer Scientific, US). Image stack alignments, scaling and adjustments were performed with Avizo software (Thermo Fisher Scientific, US) and the brightness/contrast was adjusted with ImageJ. Image processing, rendering and animations were generated with Avizo software.

### Non aqueous fractionation

8-10 mg of freeze-dried storage root material were used for NAF analyses based on a modified method of Fürtauer et al. (2016). A gradient of tetrachloroethylenes and n-heptanes with a total density of 1.35 to 1.60 (1.35; 1.40; 1.45; 1.50 and 1.60) was mixed to suspend the freeze-dried samples. After sonication (1h) in the lowest density gradient, samples were filtered and fractionated consecutively by centrifugation. Resulting supernatant was transferred to new tubes and used for determination of marker enzyme activities sugar levels measured. The resulting pellet was subsequently sonicated again in the gradient at the next higher density (20 min), filtered and fractionated again via centrifugation. Determination of different activities of marker enzymes from vacuole, cytosol and plastid was performed according to Fürtauer et al. (2016). To quantify the sugars from the different fractions, samples were first taken up in 300 µl of H2O. For further analysis, 60 µl of sample was used and analyzed by NADP-coupled enzyme assay in 96-well plates via Tecan plate reader (Tecan Infinite 200, Tecan Group). The sugar levels and enzymatic activities are listed in Supplementary File 18.

## Code and data availability

All used commands and scripts are available on github (https://github.com/Division-of-Biochemistry-Publications/CASS).

Raw sequencing reads have been deposited to NCBI’s Sequence Read Archive (submission ID: SUB13973922) under BioProject ID PRJNA1045511, available at https://www.ncbi.nlm.nih.gov/sra/PRJNA1045511.

## Acknowledgments

We thank Micheala Reiser for excellent technical assistance in the lab. Sindy Gutschker, Alexander Kaier and Stephan Reinert for helpful discussions about bioinformatics and data analysis, as well as Alfred Schmiedl for IT support. We thank Michaela Fischer-Stettler and Samuel C. Zeeman for helpful discussions and input on the manuscript.

## Author contributions

D.R. performed the greenhouse trials, NSC measurements, light-microscopy, processed and evaluated the transcriptome data, and did the statistical analysis and visualization of the data. V.V, J.K. and M.K. executed electron microscopy. L.B., B.P. and E.N. did the phloem exudate and non-aqueous fractionation experiments. S.A., and A.R.F. performed metabolic measurements. D.R. and W.Z. planned the experiments. U.S. and W.Z. supervised and coordinated the work. D.R. and W.Z. wrote the manuscript with the help of all authors.

## Conflict of interest

The authors have declared that no competing interest exists.

## Funding

U.S.: received funding from the Bill and Melinda Gates Foundation through the grant INV-008053 (Cassava Source-Sink).

